# Multiple-Context Training Enhances Generalization of Cued Fear Extinction and Requires the Dorsal Hippocampus

**DOI:** 10.64898/2026.01.18.700199

**Authors:** S Penna, S Gonzalez, E Hammerman, C Roebuck, S Patel, E Lock, C Desmond, Y Fukunaga, G Maloy, V Cazares

## Abstract

**BACKGROUND:** Multiple-session fear extinction (FE) models exposure therapy, which also occurs over many sessions. In humans, conducting extinction training across multiple contexts improves fear suppression as compared to a single context, but rodent work delineating boundary conditions and neural mechanisms remains sparse.

**METHODS:** We compared single-context FE (SCFE) versus multiple-context FE (MCFE) in 129S1 and C57BL/6J mice, manipulating context familiarity (familiar vs novel FE contexts). Outcomes included FE acquisition and tests for fear relapse such as recent (2 d) and remote (30 d) recall in trained, extinction, and novel contexts, as well as reinstatement tests. We mapped extinction-related neural activity with dual Fos labeling (TRAP×Fos immunofluorescence) and graph analyses of regional co-activation, and used chemogenetic inhibition to selectively silence dorsal (dHP) or ventral hippocampus (vHP) during extinction.

**RESULTS:** MCFE produced greater across-session reduction in freezing than SCFE. The MCFE advantage did not require contextual novelty. MCFE also enhanced short- (2 d) and long-term (30 d) recall in a novel context, but did not prevent recovery over time or US-induced reinstatement. Dual-labeling revealed that MCFE strengthened hippocampal–prefrontal co-activation (with relatively weaker prefrontal–amygdala coupling than SCFE). Chemogenetic silencing of dHP, but not vHP, selectively impaired between-session extinction under MCFE and abolished the MCFE benefit; hippocampal silencing did not improve SCFE.

**CONCLUSIONS:** Conducting extinction across multiple contexts enhances acquisition and recall relative to SCFE, without requiring novelty, and engages distinct hippocampal–prefrontal circuit dynamics. dHP activity is necessary for the MCFE-specific improvement, highlighting a circuit mechanism in which distributing extinction across contexts augments fear reduction.

## Introduction

Exposure-based therapy, a core component of cognitive-behavioral therapy, is a first-line treatment for anxiety disorders. Through repeated, safe encounters with feared cues or situations, exposure reduces maladaptive avoidance and enables fear-extinction learning. Despite its clinical value, remission rate with exposure therapy is only ∼50-55%, varying by diagnosis (1). A major contributor to these mixed outcomes is that extinction often fails to generalize beyond the treatment setting, likely explaining why up to two-third of patients relapse after 8 years (2).

Extinction reduces conditioned responding primarily through new inhibitory learning that suppresses expression of the original CS-US memory—i.e., a conditioned stimulus (CS) such as a tone no longer predicts an aversive unconditioned stimulus (US) such as a mild shock. Erasure of the initial CS-US memory also occurs, but it is limited, as evidenced by the return of fear elicited by context renewal (exposure to CS outside of the extinction training context), US-reinstatement (re-exposure to the US), and spontaneous recovery over time (3). Importantly, extinction mechanisms depend on the training regimen: in rodents, single-session extinction relies predominantly on inhibitory learning, whereas multiple-session protocols can additionally recruit erasure-like processes (4). Critically, deficits in this inhibitory learning are present in people with sustained anxiety, while stronger extinction recall predicts efficacious CBT outcomes, underscoring the need to understand how extinction generalizes across contexts (5,6).

Most preclinical studies emphasize single-session extinction, whereas clinical exposure typically unfolds over multiple sessions. Clinical guidance further recommends varying internal, external and interpersonal contexts to enhance retrievability and generalization of extinction learning (7). Although findings from laboratory studies in humans using multiple-session and multiple-context are mixed across tasks and diagnosis, they motivate further mechanistic study in rodents (8–13). Further, the neurobiological mechanisms and the conditions under which multiple-session, multiple-context extinction is most effective are not well defined in rodents.

In the present studies, we test how distributing extinction across multiple distinct contexts shapes fear reduction and its neural substrates, relative to single-context fear extinction (SCFE). We show that multiple-context fear extinction (MCFE) accelerates between-session extinction and does not require contextual novelty during training to confer its benefits. Critically, these behavioral effects generalize across two genetically distinct strains that differ in stress susceptibility (C57BL/6J and 129S1). Furthermore, we found that relative to SCFE, MCFE improves recall in novel contexts at recent and remote timepoints. At the systems level, MCFE is associated with a reorganization of hippocampal–prefrontal engrams, and dorsal hippocampal activity is necessary for the MCFE-specific enhancement during extinction.

## Methods and Materials

### Mice

Founder mice were obtained from Jackson Laboratories and bred in our colony. Naive male and female 129S1/SvImJ (Stock #002448), C57BL/6J (Stock #000664), Ai16 (Stock #007914), and Fos-CRE-ER2 (Stock #020940) were used in the experiments. All mice were housed by sex in cages of 2-5 subjects with 24/7 access to food and water *ad libitum*. Mice were on a 12 hr on/off light cycle and all experiments were conducted during the light-on phase. Prior to fear conditioning, mice were randomly assigned to SCFE (single context fear extinction) or MCFE (multiple context fear extinction) groups of about the same size with roughly equal numbers of males and females in each group. Exact numbers of mice are provided in figure captions. All procedures performed using mice were approved by and carried out in accordance with the Williams College Institutional Care and Use Committee (IACUC).

### Fear conditioning apparatus

Fear conditioning was conducted in 9.5 x 12 x 8.25’’ chambers (#VFC2-USB-M, Med-Associates, Fairfax, Vermont) housed in 24.25” x 22” x 28.75” sound attenuated boxes. Each chamber has clear acrylic backs and doors, aluminum sides, stainless steel rod floors (spaced 0.31’’ apart), stainless steel drop pans, and is illuminated by overhead near infrared (940 nm) light. Foot shocks were administered through the rods via solid state shock scramblers. Behavior was recorded at a frame rate of 30 Hz using a monochrome front-facing camera equipped with a near-infrared pass filter. *VideoFreeze* (Med-Associates) running on a desktop PC was used to acquire images and to program delivery of all stimuli.

### Cued fear conditioning

Mice were fear-conditioned once per day over the course of two consecutive days. The protocol consisted of three tones (4 kHz), twenty five seconds in length used as conditioned stimuli (CS), which were immediately followed by a two second 0.7-0.75 mA footshock (unconditioned stimulus, US). The protocol length totaled 9 minutes and 14 seconds. Both days of conditioning occurred in context A, composed of the chamber described above, no white light and a vinegar solution (30% distilled white vinegar in water) sprayed into the waste tray, producing an ambient scent. Once the protocol terminated, the mice were removed from the chambers and placed into their home cages. In between cohorts, the chambers were wiped down with the same solutions used to produce the ambient scent.

### Fear extinction, recall tests, and fear reinstatement

After fear conditioning, mice underwent up to 3 consecutive days of fear extinction consisting of 12 unreinforced CS-tones with inter-stimulus intervals ranging from 5 to 22s. For mice undergoing single context fear extinction (SCFE), each training session over the 3 days was conducted in context B, while for mice undergoing multiple context fear extinction (MCFE), each training session was conducted in a unique context (B-D). Context B was assembled in the chambers with a white round wall, a white acrylic floor, and an ethanol scent. Context C was assembled using a black triangular wall, a black acrylic floor, and a vanilla scent. Finally, context D was assembled using a green and white striped wall, a blue rubber floor mat, and a lavender scent. Every fear extinction context had overhead lights on. All scents were produced by diluting essential oils in 70% ethanol. Recall tests were carried out 2 and 30 days following fear extinction and consisted of the presentation of 4 unreinforced CS-tones. For fear reinstatement experiments, mice were re-exposed to three US-footshocks in Context A following fear extinction training.

### Chemogenetics

Mice underwent stereotaxic surgery to infuse purified adeno-associated virus (AAV) to transduce an inhibitory designer receptor activated by designer drug (AAV5-CaMKII-hM4D(Gi)-IRES-mCitrine, “Gi”), or a control fluorescence marker (AAV5-CaMKII-mCherry, “CON”). These procedures were performed 2-4 weeks prior to behavioral experimentation. In brief, mice were maintained under anesthesia with 1-3% isoflurane, secured in a stereotaxic frame and craniotomies were performed to inject AAV into dorsal (AP -1.9; ML +/- 1.5; DV -1.8; 500nl) or ventral (AP -3.16; ML +/-3.0; DV -3.85, -3.5, -3.25; 200nl/site) hippocampus. After the surgery, mice were injected subcutaneously with the analgesic carprofen (5 mg kg−1) in sterile saline (0.9%). All mice, independent of AAV condition, were habituated to handling and scruffing for 7 days prior to the start of behavioral experimentation. On extinction session day 2, 30 minutes before the start of training, mice were injected with DREADD agonist Compound 21 (C21, 1 mg kg−1), which does not undergo back-metabolism to clozapine (14).

### Dual labeling: Fos neuronal tagging and immunofluorescence

Dual fos labeling of active cell populations was achieved by combining the TRAP2 system (Targeted Recombination of Active Populations 2) and Fos immunolabeling (15). Recombination was induced via injection with tamoxifen (B5965-5000, APExBIO). 30 mg/ml TAM was dissolved directly in corn oil at 50 °C and filtered through a sterile syringe filter (0.2 μm PES, Millipore Sigma). To minimize non-specific IEG activity, mice were maintained in a holding room adjacent to the animal testing room and completely undisturbed for a full 24 hour period surrounding the TAM injection. Fear learning-related activity-dependent neuronal tagging was induced by a single intraperitoneal injection of TAM (50 mg kg−1) administered immediately after the cued fear conditioning session on day 2. To account for non-learning related neuronal tagging, one group of mice underwent the same behavioral procedures as the experimental groups (ie tone and context exposure) but did not receive footshock pairings. To label active cell populations related to extinction learning, mice were deeply anesthetized with avertin (500 mg kg−1) and transcardially perfused with 1× PBS, followed by 4% paraformaldehyde in 1× PBS. Extracted brains were post-fixed overnight at 4 °C in 4% paraformaldehyde. For immunohistochemistry, 100-μm coronal sections were collected on a vibratome, washed in 1× PBS and blocked at room temperature for 1 h in 10% normal goat serum (NDS) in 1× PBS with 0.3% Triton X-100 (PBST). Sections were incubated with primary antibodies—1:1,000 rabbit anti-Fos (Cell Signaling #2250), for 48h at 4 °C. Sections were rinsed in 1× PBST and incubated in secondary antibodies (1:1000 goat anti-rabbit Alexa 647) in 1× PBST for 1 h at room temperature. Sections were rinsed in 1× PBST and counterstained with DAPI (3uM, Invitrogen, #D1306), mounted onto slides, and coverslipped with antifade reagent in glycerol (90% Glycerol, 0.5% N-propyl gallate, 20mM Tris, pH 8.0)

### Imaging and quantification

Fluorescence images were acquired using an Olympus VS200 slide-scanning epifluorescence microscope equipped with a 10x objective (UPLXAPO, NA 0.40), an EM-CCD camera (Orca Fusion), LED illumination (X-Cite Xylis) and optical filters. Optically stitched images of brain sections were produced for TRAP (tdTomato; filters, Ex 605/70; BS 565; Em 545/25) and Fos labeling (647 immunofluorescence; filters:Ex 635/18, BS652, Em 680/42). To align imaged brains to the Allen Mouse Brain Common Coordinate Framework sections were adjusted for anterior-posterior position, angle, rotation, scale and using a deep neural network (DeepSlice, (16)). Detection of positive TRAP or Fos labeling was performed using StarDist, an automated deep-learning-based method via the QuPath software environment (17).

### Statistical analyses

Behavioral outcomes were analyzed with linear mixed-effects models that included a random intercept for subject (Animal) and fixed factors matched to each design (e.g., Group, CS, Day, and their interactions). Pre- and post-CS periods were excluded. Categorical predictors were effects-coded (sum-to-zero) so that main effects represent deviations from the grand mean; ordered factors (e.g., CS position, Day) were additionally examined for trend components (linear, quadratic) where appropriate. Omnibus (Type III) tests were used to evaluate main effects and interactions, with Satterthwaite-approximated degrees of freedom. When interactions were significant, we conducted simple-effects tests and reported Holm-adjusted pairwise contrasts; main-effect post hocs were only performed when no higher-order interaction involving that factor was significant. For visualization, we plotted estimated marginal means (EMMs) with standard errors (SE). Unless noted, tests were two-tailed with α = .05. Where reported, the standardized effect size g denotes the specified model contrast divided by the model’s residual standard deviation.

## Results

### Extinction training in multiple contexts enhances memory savings

We first examined the effects of FE training in single vs. multiple contexts on cued fear learning and memory using two mouse strains: the stress-susceptible 129S1 (S1; previously reported to exhibit deficits in FE) and the more stress-resilient strain, C57BL/6J (B6). Mice underwent two sessions of fear conditioning, each consisting of 3CS-3US pairs (4kHz tone, 0.7-0.75mA footshock). Following fear conditioning, animals received three sessions of FE training conducted either in the same context (B,B,B; SCFE) or three distinct novel contexts (B,C,D; MCFE). Each extinction session consisted of 12 unreinforced CS presentations with sessions spaced 24h apart (experimental timeline, **Fig. 1A**).

**Figure 1.**
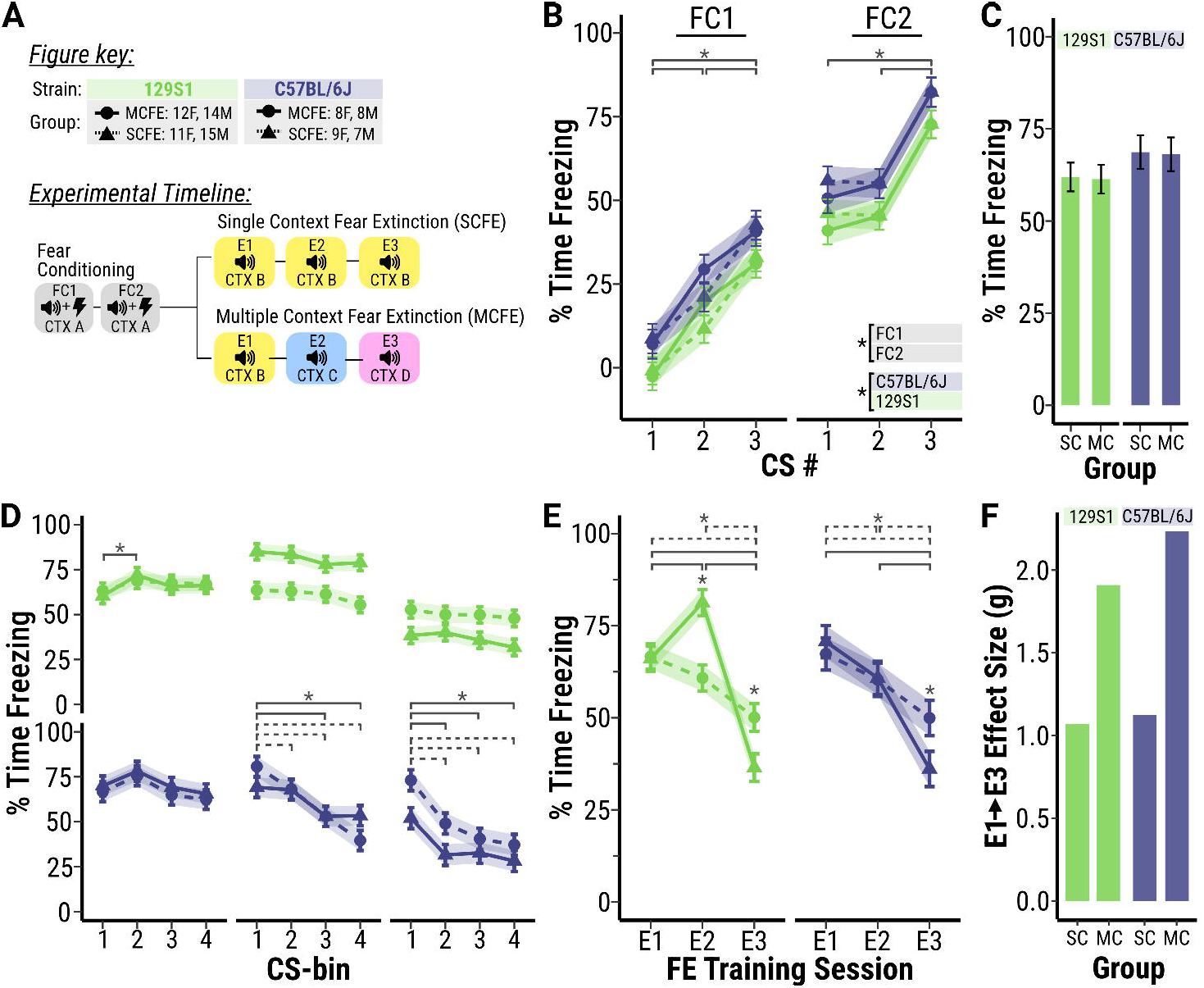
Extinction training in multiple contexts enhances memory savings. **(A)** Experimental timeline and figure key. **(B)** Cued fear conditioning (FC). Three CS-US pairings on consecutive training days (FC1, FC2) increased conditioned freezing in both C57BL/6J (B6) and 129S1 (S1) mice from CS1 to CS3. **(C)** Fear recall at the start of extinction (E1). Freezing to the first 3 CS did not differ between groups, confirming comparable conditioned fear before extinction. (**D-E)** Extinction sessions (E1–E3) were conducted in either a single context (SCFE) or multiple contexts (MCFE). **(D)** Within-session extinction. Freezing was averaged in three-CS bins and compared within each session. Only B6 mice showed significant within-session decreases on E2 and E3, independent of extinction type. **(E)** Between-session extinction all groups exhibited significant reductions in freezing across extinction days (E1→ E3). MCFE mice exhibited lower levels of freezing on E3, with S1 MCFE animals showing a transient increase in freezing in E2 (context shift B → C), consistent with renewal, followed by a sharp decline on E3. **(F)** Standardized effect size (g) for the E1→E3 reduction in freezing. MCFE mice showed larger across-session decreases than SCFE mice. g = a standardized mean difference, computed as the E1–E3 contrast divided by the model residual SD. Shaded areas show EMM ± SEM. Sample sizes (panel A). Asterisk denotes significant effects (P <0.05) from a Type III ANOVA with Holm-adjusted contrasts.

Freezing during fear conditioning was modeled as a function of Group (SCFE, MCFE), CS (tones 1–3), and conditioning day (FC1, FC2), including all two- and three-way interactions, with Sex and Strain (B6, S1) entered as covariates. Because the SCFE and MCFE groups experienced identical procedures during conditioning, we expected no group differences at this stage. ANOVA revealed significant main effects of CS (F[2,410] = 71.5, p < 0.001), conditioning day (F[2,410] = 307.26, p < 0.001), and strain (F[1,80] = 11.25, p < 0.001), as well as a CS × day interaction (F[2,410] = 4.41, p < 0.01; **Fig. 1B**). As predicted, there was no effect of Group (p = 0.967) and no effect of Sex (p = 0.462). Post-hoc contrasts confirmed increased freezing across CS presentations within each day (FC1: CS1 < CS2 < CS3; FC2: CS1 < CS3 and CS2 < CS3; all p < 0.05). Thus, fear acquisition increased across trials and days, with no differences between experimental groups or sexes.

To confirm that both groups began extinction from comparable levels of conditioned fear, we next compared group freezing during the first three CS of the initial extinction session (context B for all animals). Freezing to the first three CS presentations—when minimal extinction learning is expected—did not differ between groups (**Fig. 1C**), indicating that SCFE and MCFE mice started extinction training with equivalent fear recall.

Freezing during extinction was analyzed as a function of Group (SCFE, MCFE) x FE training day (E1-E3) x and CS-bin (1-4, composed of 3CS blocks per bin), with Strain (B6, S1) and Sex included as covariates. The model captures both within-session (**Fig. 1D**) and across-session (**Fig. 1E**) dynamics of fear reduction. The ANOVA revealed a main effect of CS-bin F[3,720] = 29.65, p < 0.001) and FE-day (F[2,80] = 64.71, p < 0.001), indicating significant decreases in freezing within and across extinction sessions. We also discovered several 2-way and 3-way interactions summarized below.

At the within-session level (**Fig. 1D**), freezing declined significantly across CS-bins in B6 mice, particularly during later extinction sessions (E2-E3). For example, in B6 SCFE animals, freezing on E3 decreased from Bin 1 to Bins 2–4 (all p < 0.001) and a similar pattern was observed in B6 MCFE mice (Bin 1 > Bins 2–4 (all p < 0.001). In contrast, S1 mice showed little to no within-session reductions, consistent with previous reports (18). At the between-session level (**Fig. 1E**), all groups exhibited progressive reductions in freezing as a function of FE Day (F[2,80]= 64.71, p < 0.001). However, a significant Group x Day interaction (F[2,80] = 11.35, p < 0.001) indicated that reductions were greater in the MCFE condition. Post-hoc contrast confirmed that by E3, MCFE mice froze significantly less than the SCFE mice of both strains (S1: *t*_79_ = 2.57, *p* = 0.012; B6: *t*_79_ = 2.04, *p* = 0.044). Additionally, MCFE produced markedly larger reductions in freezing than SCFE across days, with large standardized effects from E1–E3 (S1 MCFE: *g* = 1.63, p < 0.001; B6 MCFE: *g* = 1.50, p < 0.001), whereas corresponding reductions under SCFE were smaller (S1: *g* = 0.91, p < 0.001; B6: *g* = 0.74, p < 0.01).

Although between-session extinction was enhanced under MCFE, this pattern was not entirely monotonic. S1 MCFE mice displayed a transient increase in freezing between E1 and E2 (t[80] = −3.42, p < 0.001) when the extinction context shifted from B to C, consistent with a renewal of conditioned fear. By E3, however, freezing levels declined sharply, indicating that extinction overcame this renewal effect. Together, these results demonstrate that extinction training produced robust within- and across-session decreases in conditioned freezing, that extinction was enhanced by the end of training in the MCFE conditions, and that extinction learning was generally slower and more context-dependent in the stress-susceptible S1 strain.

### Multiple novel or familiar contexts enhance extinction memory

Novel experiences have been shown to promote encoding and persistence of memory traces at physiological and behavioral levels (19–21). To test whether contextual novelty was necessary to enhance fear extinction memory, we compared single-context extinction training (SCFE) to training in multiple *familiar* (F-MCFE) and multiple *novel* contexts (N-MCFE) using the 129S1 strain. The procedure followed Figure 1 except that 24h before training a 20 minute pre-exposure was carried out to either: the subsequent extinction training context (F-MCFE; familiarizing the FE context), an irrelevant context (N-MCFE; leaving the FE contexts novel at training), or the same context (SCFE; context B). Therefore, animals in the N-MCFE group encountered novelty during FE, whereas F-MCFE animals extinguished in distinct but familiar contexts (see **Fig 2A**).

**Figure 2.**
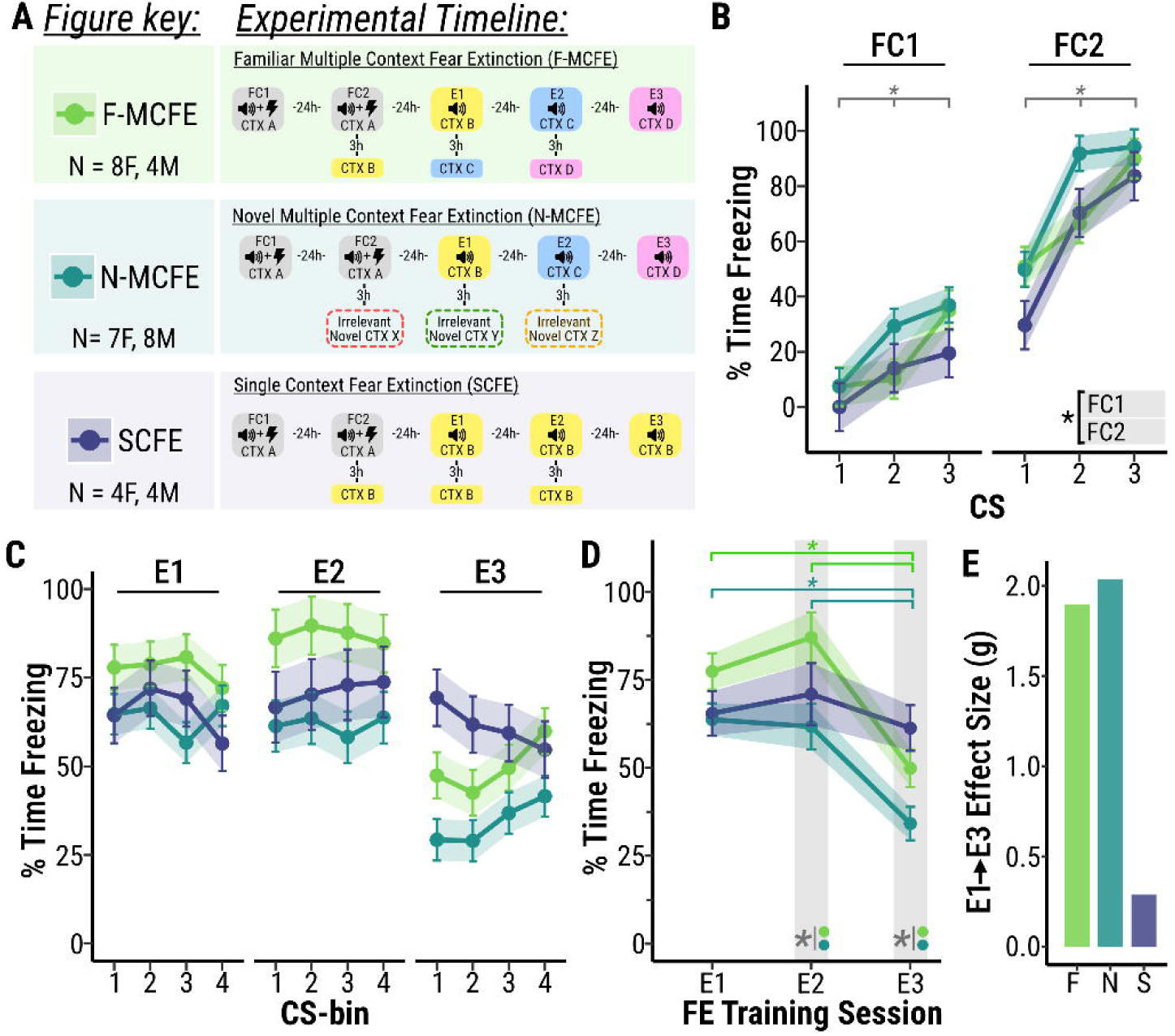
Multiple familiar or novel contexts enhance extinction memory. **(A)** Experimental timeline and figure key. Procedures matched those in Fig. 1 except for the pre-exposure 24 h before extinction: F-MCFE mice were pre-exposed to the subsequent extinction contexts, N-MCFE mice to an irrelevant context (leaving extinction context novel), and SCFE mice to context B. **(B)** Fear conditioning (FC). CS-US pairings on consecutive training days (FC1, FC2) increased conditioned freezing across CS and FC days irrespective of extinction groups. **(C–D)** Fear extinction (E1–E3) were conducted in either a single context (SCFE) or multiple contexts (MCFE). **(C)** Within-session extinction. Freezing (averaged in three-CS bins) showed no significant within-session decreases. **(D)** Between-session extinction. Both N-MCFE and F-MCFE reduced freezing from E1→E3 and E2→E3, whereas SCFE showed no across-day reduction. On E2, N-MCFE froze less than F-MCFE; on E3, N-MCFE froze less than both F-MCFE and SCFE. **(E)** Standardized effect size (g) for the E1→E3 reduction. Both MCFE conditions showed larger across-session decreases than SCFE. Conventions as in Fig. 1; g and SEM as defined there.

Analysis of FC data (**Fig. 2B**) revealed a significant conditioning day x CS-bin interaction (F[2, 160]=4.83, p < 0.001), with no main or interaction effects of FE conditions (SCFE, F-MCFE, N-MCFE). Follow-up contrast confirmed that all animals showed higher freezing across CS-bin presentations within each FC day (p<0.01), as well as greater freezing on FC2 compared to FC1(p<0.0001) indicating robust fear acquisition.

Results from the FE data analysis (**Fig. 2C-D**) revealed a significant Extinction Group x FE-Day interaction (F[4,34.99] = 4.19, p < 0.01). Within-group contrasts showed that both F-MCFE, N-MCFE groups reduced conditioned freezing in E3 relative to E1 and E2 (p<0.001), whereas SCFE showed no significant change across extinction sessions. Between group comparisons on each FE-Day indicated that the F-MCFE and N-MCFE groups differed on E2 (p=0.0375) and E3 (p<0.01). Standardized E1–E3 effect sizes were large for both multiple-context conditions (F-MCFE: *g* = 1.9, p < 0.01; N-MCFE: *g* = 2.03, p < 0.0001) and small/non-significant for SCFE (*g* = 0.29, p = 0.697). Together, these data indicate that contextual novelty is not required to enhance extinction with multiple-context training relative to SCFE, though novelty may confer an additional benefit particularly later in training.

### Extinction training in multiple contexts limits the return of fear in novel contexts

Extinction training does not erase the original CS-US memory, leaving animals susceptible to the return of fear via spontaneous recovery (with passage of time), renewal (in new contexts), or reinstatement (with US re-exposure) (3). We asked, then, whether MCFE confers greater resistance to fear relapse relative to SCFE.

To assess time- and context-dependent relapse, three cohorts of 129S1 mice that underwent SCFE and MCFE training were tested 2 and 30 days after extinction. Cohort 1 was tested in the original fear conditioning context, cohort 2 was tested in two novel contexts (E and F), and cohort 3 was tested first in the extinction context (B) and subsequently in a new context (E, see the top of **Fig 3A**).

**Figure 3.**
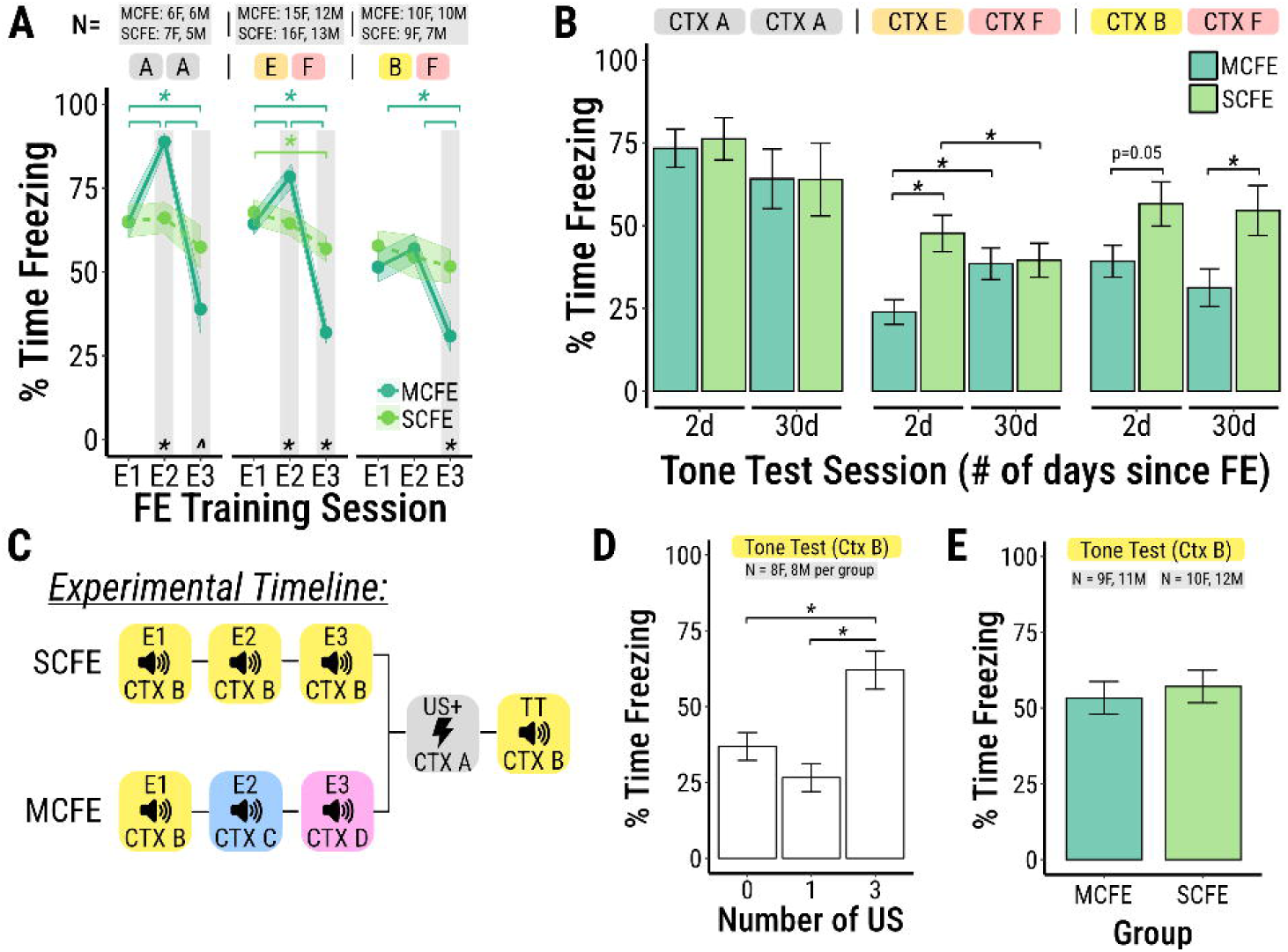
Extinction training in multiple contexts limits the return of fear. **(A)** Procedures matched Fig. 1. As before, groups were treated identically up to extinction, and MCFE mice showed lower freezing than SCFE by E3 (see Figs. 1–2). **(B)** Recall was tested at 2 days (recent) and 30 days (remote) in three cohorts: testing in A (conditioning context), in novel contexts (E,F), or extinction context B at 2 d, novel context F at 30 d. Left: No group differences in context A at either timepoint. Middle: In novel contexts, MCFE < SCFE at 2 d, but the group difference was absent at 30 d; MCFE increased freezing from 2 to 30 d. Right (B,F): A group main effect was observed; MCFE tended to freeze less at 2 d in B and showed lower freezing than SCFE at 30 d in F. **(C)** Reinstatement timeline. **(D)** Unsignaled US exposure in context A followed by testing in B: three footshocks—but not one—reinstated freezing. **(E)** After 3-US reinstatement, MCFE and SCFE showed comparable freezing. Conventions as in Fig. 1.

As in previous experiments (**Fig. 1** and **Fig. 2**) there were no differences in fear acquisition between extinction groups (SCFE vs. MCFE) in any cohort, and the MCFE enhancement in between-session extinction was replicated (**Fig. 3A**). We then assessed differences in 2d and 30d recall across the three cohort designs: In cohort 1, there were no differences between mice tested in the original fear conditioning context (A) at either timepoint. In cohort 2 (two novel contexts E and F), a significant Extinction Group (SCFE, MCFE) x Test Day (2d, 30d) interaction emerged (F[1,377.68] = 34.26, p < 0.001). Post-hoc tests revealed that SCFE and MCFE groups differed at the 2d (t[84.06]=3.94), but not the 30d (t[101.26]=-0.86) tone-test.

Within groups, SCFE mice decreased freezing from 2 and 30d tone-tests (t[382.18]=4.18, p < 0.001), consistent with additional fear extinction; whereas the MCFE mice increased freezing (t[378.73]=-4.05, p < 0.001), consistent with fear recovery. To test whether SCFE reductions in freezing between 2d and 30d tone-tests reflected exposure to two novel contexts (a pseudo-MCFE effect), cohort 3 was designed such that animals were tested in the familiar context B (2d), followed by novel context F (30d). This cohort showed a main effect of the extinction group (F[1,34]= 7.85, p < 0.01), with no test day effect or interaction. We examined the Group effect at the 30-d test in the novel context and found that MCFE froze less than SCFE (t[58.7]=2.69, p < 0.01). These results suggest that lower freezing levels in the cohort 2 SCFE compared to MCFE were due to a pseudo-MCFE effect. Overall, these findings support a persistent MCFE advantage at the long-term test in a novel context. Across designs, MCFE enhanced extinction recall in novel contexts at 2 days (Cohort 2, E) and remained lower than SCFE at 30 days in a novel context (Cohort 3, F).

To compare the sensitivity of SCFE and MCFE susceptibility to fear reinstatement, we measured freezing after unsignaled US-re-exposure. We first established a reinstatement threshold, finding that 3 footshocks–but not 1–were sufficient to reinstate freezing (t[33]=-3.42, p < 0.01), **Fig 3D**). Using this 3-US protocol, SCFE and MCFE mice did not differ in freezing levels during a tone test 24h later (p> 0.05). Thus, MCFE limits context-dependent relapse (renewal) at recent and remote timepoints, but may not prevent relapse driven by time (spontaneous recovery) or US re-exposure (reinstatement).

### The number of sessions and contextual change during FE alters neural activation in key brain regions

To identify brain regions engaged by MCFE, we quantified neural activation during FC and FE using a dual Fos mapping strategy. Mice underwent SCFE or MCFE as in **Fig. 1**, with two additional groups: SSFE (single session FE) and US− (context/tone exposure without shock, indexing background Fos). Putative conditioning engrams were tagged with TRAP2 during FC, and extinction-related activation was assayed with Fos immunohistochemistry (Fos-IF) after FE. This TRAP2×Fos-IF approach enabled comparison of the magnitude and regional distribution of FC- and FE-related populations and their overlap.

For FC analyses, we observed significant main effects of Group (F[3,79.88]= 10.77, p < 0.001), CS (F[2,298.78]= 34.4, p < 0.001), and Day (F[1,337.3]= 245.91, p < 0.001), with Group × CS (F[6,298.78]= 2.73, p < 0.05) and Group × Day (F[3,336.75]= 18.81, p < 0.001) interactions. Post-hoc tests showed that, except for US−, all groups exhibited increasing freezing across CS presentations on FC1 and FC2. During FC2 (the TRAP-labeling session), all FE groups (SSFE, SCFE, MCFE) froze more at each CS than the US− group (p < 0.05).

Analysis of behavior during FE (**Fig. 4B**) yielded main effects of Group (F[3,117.09]= 16.46, p < 0.001) and Day (F[2,815.39]= 5.27, p < 0.01), as well as Group × CS-bin (F[9,812.87]= 2.97, p < 0.01) and Group × Day (F[6,815.11]= 17.68, p < 0.001) interactions. Critically, these data replicate earlier results (**Figs. 1–3**): MCFE mice showed lower freezing than SCFE by E3. For SSFE, mice were placed in the conditioning chamber, freezing was recorded across all sessions, but CSs were presented only on E3; as expected, SSFE displayed significantly higher freezing on E3 than on E1–E2 (where no CSs were presented) and higher E3 freezing than both SCFE and MCFE. The US− group exhibited lower freezing than all other groups across FE sessions (p < 0.05).

**Figure 4.**
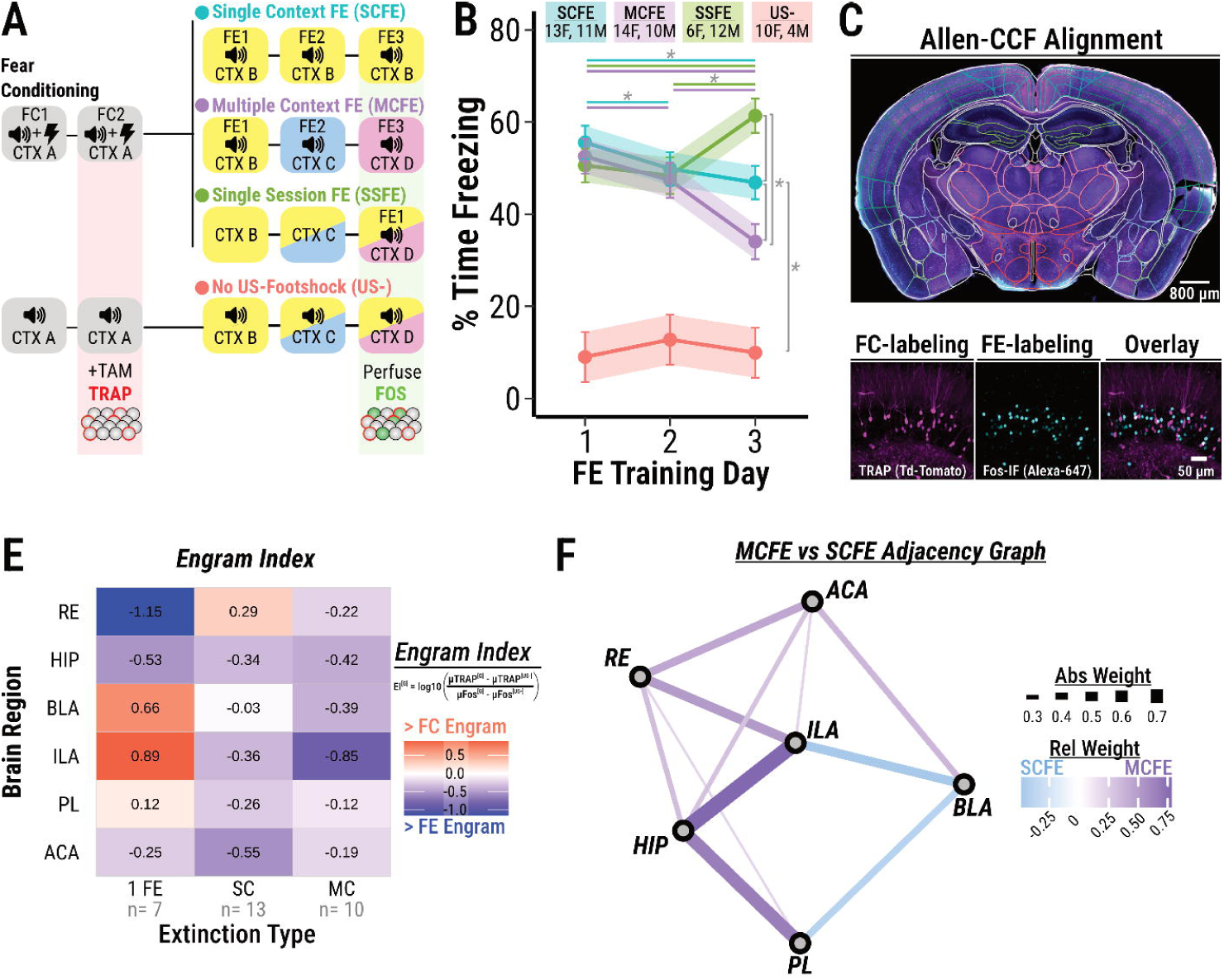
MCFE results in greater extinction-related activity and hippocampal-prefrontal co-activation. **(A)** Experimental timeline. As in Fig. 1, with two exceptions: (1) two additional groups were included—US− (context + tone exposure; no shocks) and single session FE (SSFE); (2) all mice received tamoxifen on FC2 to TRAP activity and were perfused 90 min after E3 for Fos immunolabeling of extinction-related activity. **(B)** Between session extinction. **(C)** Representative section and Fos-labeling. **(D)** Engram index (EI-G). The magnitude of conditioning-engram change versus extinction-engram change (each relative to the US− control). EI-G; values < 1 indicate greater extinction-evoked than conditioning-evoked activation. The heatmap summarizes regional EI-G across groups. **(E)** Difference graph (MCFE − SCFE). Graph derived from group-level adjacency matrices of regional TRAP⁺ ∩ Fos⁺ densities. Each edge reflects the difference in pairwise correlation (Δr) between regions (MCFE vs. SCFE); purple edges indicate stronger co-activation in MCFE, blue stronger in SCFE; edge thickness Δr. The pattern suggests distributed reorganization of the extinction engram, with relatively stronger hippocampal–prefrontal–thalamic co-activation under MCFE.

To focus on behaviorally relevant regions, we first restricted analyses to areas with greater TRAP⁺ (FC tagging) and Fos⁺ (FE tagging) counts than US− controls [one-sided tests with multiple-comparison control, e.g., FDR q<0.05 across regions]. We then computed an engram index (EI^G^,**Fig. 4E**) to identify the relative expression of the putative FC engram vs. FE engrams (values <0 favor FE over FC). We found that relative to SSFE and SCFE, MCFE showed a larger shift toward FE (lower EI^G^) in hippocampus (HP), infralimbic cortex (IL), and basolateral amygdala (BLA). Across the surveyed regions, MCFE tended to show low EI^G^ (greater FE labeling), whereas SSFE favored FC labeling in prelimbic cortex (PL)/IL/BLA and SCFE favored FC labeling in reuniens (RE).

To probe functional circuit organization, we examined pairwise regional co-activation (correlations of dual-labeled densities) and visualized the MCFE–SCFE difference graph (**Fig. 4F**). MCFE showed stronger HP↔medial prefrontal cortext (mPFC: ACA/PL/ILA) co-activation, whereas SCFE showed stronger mPFC (PL/IL)↔BLA coupling. These data suggest that MCFE redistributes engram-related activity from prefrontal–amygdala toward hippocampal–prefrontal pathways, implicating HP and mPFC as hubs for context-general extinction.

### Chemogenetic inhibition of the dorsal, but not ventral, hippocampus impairs fear extinction training in multiple contexts, while leaving single-context extinction intact

The hippocampus is essential for contextual retrieval, allowing organisms to respond to fear cues in a manner appropriate to the surrounding context (22). We asked whether enhanced extinction retrieval in MCFE depends specifically on the dorsal hippocampus, which is rich in place cells, and/or the ventral hippocampus, which contains cells responsive to stimulus identity and intensity (23). To test this, we compared mice expressing inhibitory chemogenetic receptors (hM4DGi, GI) with those expressing a control reporter (mCherry, CON) in either the dorsal (**Fig. 5 A-B**) or ventral hippocampus (**Fig. 5 C-D**) during MCFE or SCFE.

**Figure 5.**
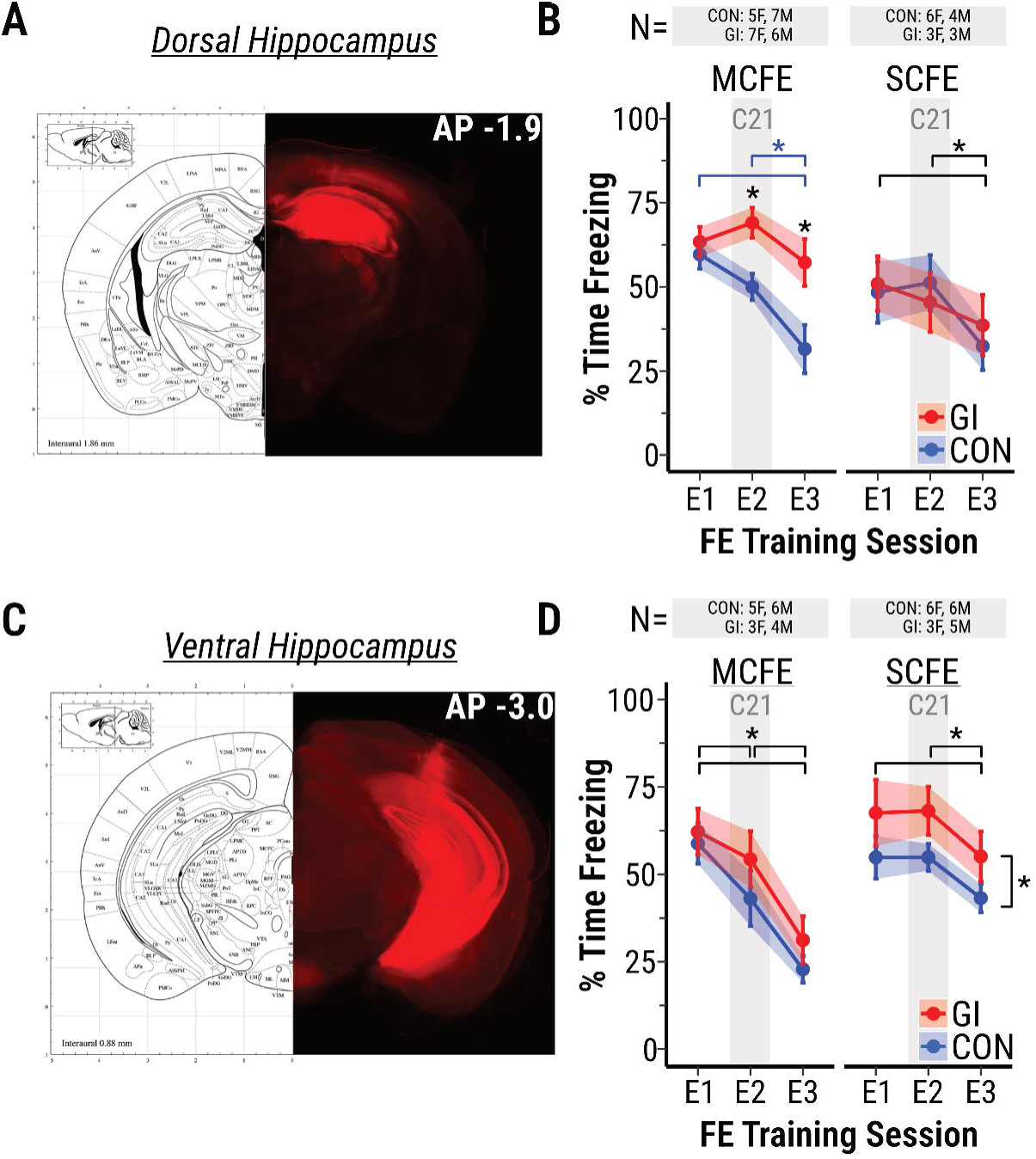
Chemogenetic inhibition of the dorsal but not ventral hippocampus disrupts extinction in multiple-contexts. **(A)** Dorsal hippocampal (dHP) viral transduction. Mice received bilateral injections of AAV5-CaMKII-hM4D(Gi)-IRES-mCitrine (Gi) or AAV5-CaMKII-mCherry (CON). Composite overlays of fluorescence images for all Gi mice. **(B)** Between-session extinction, dHP. Fear conditioning (data not shown) and fear extinction (E1–E3) were as in Fig. 1, except all mice received the DREADD agonist 21 (C21; 1 mg kg⁻¹ i.p.) 30 min before E2. In the multiple-context fear-extinction (MCFE) cohort, Gi mice froze more than CON mice on E2 and E3 and failed to reduce freezing from E1 to E3. In the single-context fear-extinction (SCFE) cohort, Gi and CON groups did not differ. **(C)** Ventral hippocampal (vHP) viral transduction. Composite overlays of fluorescence images for all Gi mice. **(D)** Between-session extinction, vHP. C21 produced no virus-specific effect in MCFE. We observed a main effect of virus conditions in SCFE wherein GI mice exhibit higher freezing than CON mice. Conventions as in Fig. 1.

After a minimum of two weeks for viral expression and recovery (**Fig. 5A, 5C**) mice underwent FC and FE as described in **Fig. 1**, with one key modification: all animals received an intraperitoneal injection of C21 (1 mg/kg in saline) 30 minutes before E2, the session in which MCFE mice first encounter the CS in a distinct context. In MCFE mice, GI expression in dHP–but not vHP– resulted in greater freezing on E2 (t[35.65]=-2.84, p < 0.001) and E3 (t[35.65]=-3.84, p < 0.0001) and abolished between session extinction (i.e. no E1-E3 reduction). In SCFE mice, neither dHP nor vHP GI expression affected between-session extinction relative to controls; however, GI-expressing SCFE mice exhibited higher overall freezing than SCFE controls (t[18]=-2.25, p < 0.05), suggesting possible anxiogenic effects of general vHP inhibition. Overall, these data suggest that silencing the dHP disrupts between-session learning in MCFE, whereas vHP inhibition leaves between-session extinction intact in both extinction regimens.

## Discussion

A major source of relapse in fear- and stress-related disorders stems from the failure of extinction memories to generalize across distinct or novel contexts. In rodents, neural ensembles encoding learned fear are more likely to be reactivated outside of the extinction context, while extinction memories remain tightly bound to the training environment (22,24). In humans, multivariate fMRI studies similarly show that activity patterns for aversive memories generalize across contexts and types of threat, whereas extinction produces new context-specific representations (25–27). Together, these findings suggest that extinction often creates new, context-bound representations rather than modifying the original fear memory— a limitation we sought to overcome with MCFE.

In the present study, we found that across two genetically distinct strains (C57BL/6J and 129S1) with disparate stress susceptibilities, MCFE accelerated between-session extinction and did not require contextual novelty during training to be effective. Critically, relative to SCFE, MCFE improved recall of extinction in novel contexts at recent and remote timepoints, suggesting a lasting effect on FE generalization. We further showed that MCFE reorganizes hippocampal–prefrontal engram coactivation patterns and, specifically, that dorsal hippocampal activity is necessary for the MCFE-specific enhancement of extinction learning.

From a learning-theoretic perspective, effective extinction (in laboratory and clinical settings) depends on prediction error, the violation of expectations of threat (28,29). In SCFE, repeated CS exposures within a single context progressively reduce prediction error, potentially limiting plasticity (30). This behavioral difference has clear implications for underlying circuit dynamics. For example, extinction-related responsiveness in infralimbic cortex (IL) and basal amygdala (BA) diminishes after the first session, suggesting that additional within-context trials may yield diminishing returns (4). By contrast, because MCFE takes place in a new context, where fear memories are predicted to be more highly engaged, MCFE is expected to maintain higher prediction error across sessions while still providing extensive CS exposure.

Contextual memory is important for gating which response among a repertoire of previously learned behaviors is best fitted to the situation — a process that is especially critical for extinction generalization. Key hubs gating CS representations across contexts are the mPFC and HP (31). We found that the dorsal but not ventral hippocampus is essential for the effects of MCFE. This is consistent with the role of the dorsal hippocampus in spatial/contextual discrimination, whereas the ventral hippocampus is more closely linked to affective processing and stimulus identity (23,32). Furthermore, using a Fos-mapping approach, we found that SCFE preferentially engages mPFC–BLA coactivation, whereas MCFE shifts network organization toward stronger mPFC–hippocampal coordination, suggesting mechanistic differences in extinction learning between paradigms. The circuit mechanism for the effect of MCFE remains undetermined, especially since there are very few direct projections from dHP to mPFC (33,34). Our data also suggest that circuits involving the RE are a plausible mechanism, since RE also shows enhanced coactivation in MCFE; neuroanatomically, it projects bidirectionally to both mPFC and hippocampus and has been implicated in fear relapse (35).

Human studies using electrophysiological and fMRI-based measurements show that the specificity of cue-related activity patterns in PFC and other regions of the extinction network is higher during extinction than during fear acquisition (25,27). This is consistent with the notion that extinction learning often represents an exception to the original CS–US rule established during fear learning. We present evidence that, at the behavioral level, MCFE enhances generalization of fear extinction, and at the neural level, it reorganizes extinction networks in a manner dependent on the dorsal hippocampus. These findings raise the possibility that MCFE transforms extinction from a context-specific exception into a more generalizable safety rule.

## Acknowledgments

Funding: National Institutes of Health, R15MH129947; Massachusetts Life Science Center, Workforce Development Capital Grant; Williams College, Class of ‘45 World Fellowship; Brain Behavior Research Foundation, NARSAD Young Investigator Grant

Non-author contributors/collaborators: Johanna Huarachi, Nikitah Gaju, Liriana Valerio, Celeste Cruz Rivera, Mayor Watts, Amy Zhao, Emma Schulman

